# Rapid reprogramming and stabilisation of homoeolog expression bias in hexaploid wheat biparental populations

**DOI:** 10.1101/2024.08.01.606180

**Authors:** Marek Glombik, Ramesh Arunkumar, Samuel Burrows, Sophie Louise Mogg, Xiaoming Wang, Philippa Borrill

## Abstract

- Differences in the relative level of expression of homoeologs, known as homoeolog expression bias (HEB), are widely observed in allopolyploids. While the evolution of homoeolog expression bias through hybridisation has been characterised, on shorter timescales the extent to which homoeolog expression bias is preserved or altered between generations remains elusive.
- Here we use biparental mapping populations of hexaploid wheat (*Triticum aestivum*) with a common Paragon parent to explore the inheritance of homoeolog expression bias in the F_5_ generation.
- We found that homoeolog expression bias is inherited for 26-27% of triads in both populations. Most triads (∼70%) conserved a similar homoeolog expression bias pattern as one or both parents. Inherited patterns were largely driven by changes in the expression of one homoeolog, allowing homoeolog expression bias in subsequent generations to match parental expression. Novel patterns of homoeolog expression bias occurred more frequently in the biparental population from a landrace x elite cross, than in the population with two elite parents.
- These results demonstrate that there is significant reprogramming and stabilisation of homoeolog expression bias within a small number of generations that differs significantly based on the parental lines used in the crossing.

## Introduction

Polyploidy, characterised by the presence of multiple chromosome sets due to whole genome duplication, is widespread amongst flowering plants (Leitch & Leitch, 2008). This duplication provides an important route to increase phenotypic diversity and adaptability to extreme environments (Van de Peer *et al*., 2017; Van de Peer *et al*., 2020). The enhanced adaptability may be mediated by the increased genetic variation within a polyploid genome and by buffering effects of the duplicate gene copies which may act redundantly (Van de Peer *et al*., 2009; Doyle & Coate, 2019). These genetic advantages of polyploidy have contributed to the success of many domesticated crop species (Salman-Minkov *et al*., 2016; Cheng *et al*., 2018).

Polyploid species can be formed by autopolyploidy, genome duplication within a single species, or allopolyploidy, hybridisation of two species followed by genome doubling. Allopolyploids contain gene duplicates known as homoeologs which originate from the different parental species. Homoeologs are frequently expressed at different levels from each other, resulting in homoeolog expression bias (HEB), which has been documented in multiple species including major crops such as cotton, wheat and brassica (Yoo *et al*., 2013; Chalhoub *et al*., 2014; Ramírez-González *et al*., 2018). In several allopolyploid crop species subgenome dominance occurs, with the dominant subgenome showing higher gene retention, more tandem gene duplications, higher gene expression and lower DNA methylation (Alger & Edger, 2020). In these cases, the higher expressed homoeologs from the dominant subgenome can more strongly control the observed phenotypes. For example, in maize the dominant subgenome contributes more to phenotypic variation (Renny-Byfield *et al*., 2017) and in strawberry key traits such as flavour and aroma are largely controlled by pathways encoded in the dominant subgenome (Edger *et al*., 2019). However, global subgenome dominance is not detected in other polyploid species such as *Camelina sativa* (Kagale *et al*., 2014), oat (Kamal *et al*., 2022) and wheat (IWGSC *et al*., 2018); instead HEB varies between individual genes within these polyploids. Several traits of critical agronomic importance have emerged from changes in the expression level of one homoeolog, resulting in a change in HEB. For example, in wheat, *cis*-variants causing higher expression levels of one homoeolog in the flowering genes *VRN1* and *PPD1* have been essential to adapt flowering time to suit a wide range of environmental conditions (Shaw *et al*., 2012; Shcherban & Salina, 2017). Moreover, whilst homoeolog sequences are similar, they are non-identical in allopolyploids and differences in HEB may have functional consequences through their translation into different protein sequences.

Studies comparing allopolyploids to their parental species have shown that HEB is a product of the original expression levels in parental species combined with alterations to expression during evolution (Grover *et al*., 2012). HEB in older polyploids differs from the parental species, which could be explained by millions of years passing since the whole genome duplication event (e.g. maize and soybean (Zhao *et al*., 2017)). However, HEB is also altered in evolutionarily young polyploids which have only existed for thousands of years such as *Brassica napus* (Chalhoub *et al*., 2014) and hexaploid wheat (Ramírez-González *et al*., 2018). Rapid alterations in HEB have been shown using resynthesized allopolyploids in species such as Brassica, Arabidopsis, cotton and wheat over a matter of generations (Adams *et al*., 2003; Wang *et al*., 2004; Yoo *et al*., 2013; Yang *et al*., 2016; Ramírez-González *et al*., 2018; Bird *et al*., 2021; Vasudevan *et al*., 2023) and in natural Tragopogon allotetraploids which are under 100 years old (Boatwright *et al*., 2018). As well as being rapid, the patterns of HEB may be conserved between different polyploidisation events due to the influence of parental legacy as observed in allopolyploid *Brassica napus* and wheat (Li *et al*., 2021; Banouh *et al*., 2023). For example, similar HEB patterns emerge between independent resynthesized lines in *Brassica napus* with > 70% of homoeolog expression bias being consistent (Bird *et al*., 2021). However, in *Capsella bursa-pastoris* there was a high degree of variability in HEB between resynthesized allotetraploid individuals, which could be ascribed to variations in segregation and recombination of homoeologous chromosomes (Duan *et al*., 2023).

Whilst considerable effort has been assigned to understanding the evolution of HEB and its relationships to the parental species, far less is known about within species variation in HEB. Hexaploid wheat, is an example of an allopolyploid without subgenome dominance, yet 30% of genes with three homoeologs (known as triads) show unbalanced HEB (i.e. unequal expression levels of homoeologs) (Ramírez-González *et al*., 2018). HEB varies between wheat cultivars (Ramírez-González *et al*., 2018) and variation in HEB across diverse wheat accessions is strongly influenced by *cis* regulation (He *et al*., 2022). Although, the standing variation in HEB within wheat has been explored, cultivars with different HEB are routinely combined in crossing schemes and the outcome for HEB is currently unknown. Is HEB from the parents maintained or are novel HEB patterns generated? If so, which factors influence HEB in the progeny and how variable is this within a population?

Here, we explore what happens to HEB over short time periods (five generations) which could be representative of plant breeding programmes. We examined two biparental populations, one population with elite parents and one population from a landrace x elite cross. We found that in both populations HEB is inherited for 26-27% of triads. We explored whether parental HEB patterns were conserved and to what extent novel HEB patterns were generated. We also carried out expression quantitative trait loci (eQTL) mapping to identify *cis-* and *trans-*variants altering HEB and found that some of these eQTL may be related to agriculturally important traits.

## Materials and Methods

### Plant materials and growth

All wheat seeds were obtained from the John Innes Centre Germplasm Resources Unit. These included the parental lines: Paragon (W10074 PF-1), Charger (WPxCHA – 1001) and Watkins 1190094-1 (WATDE0228) and F_5_ lines from two bi-parental mapping populations generated by single seed descent: 161 F_5_ lines for Paragon x Charger (PxC) and 50 F_5_ lines for Paragon x Watkins 1190094-1 (PxW).

For each parental and individual F_5_ line, five replicate seeds were germinated on moist filter paper in a petri dish. The dish was wrapped in aluminium foil and placed in a 4°C fridge. PxW and PxC seeds were kept in the fridge for three and five days, respectively and then kept at room temperature for 3 days. Seedlings were then transplanted into 24-well trays of Levington M2 compost utilizing a randomized block design for replicates and genotypes and grown in heated and lit greenhouses with setpoints: 20°C day, 16°C night with 16 hours light. Tray order was changed every 2-3 days to minimise microenvironmental effects (lighting, temperature, etc) biasing plant growth. PxW seedlings were sown in April 2021 and leaf tissue was harvested 16 days after transplantation. PxC seedlings were sown in June 2021 and leaf tissue was harvested 14 days after transplantation.

The second and/or third youngest leaves were sampled from each plant with 3-4 cm of leaf tissue cut and placed into a 1.5ml microcentrifuge tube, before being immediately flash frozen in liquid nitrogen and stored at −70°C. For both crosses, tissue harvesting began at 9 am and ended by 3.30 pm on the same day with 3-4 experimenters involved in the harvest.

### RNA extraction

Three biological replicates were used for RNA extraction. Leaf tissue was homogenised using the TissueLyser II (Qiagen) or Geno/Grinder 2010 (Spex) and one to two 3 mm tungsten carbine beads per tube. Total RNA was extracted from leaf tissues using the RNeasy Plant Mini Kit (Qiagen, 74904) with an optional on-column DNase digest using the RNase-Free DNase Set (Qiagen, 79254). If required, RNA was cleaned using the RNeasy RNA cleanup protocol. mRNA library preparation with polyA selection and 2×150 bp paired end sequencing on the Illumina NovaSeq was done by GENEWIZ UK Ltd.

### HEB analysis

Raw RNA-seq reads were pseudo-aligned to the IWGSC Chinese Spring transcriptome reference v1.1 using kallisto ver. 0.46.1 (Bray *et al*., 2016; IWGSC *et al*., 2018) with default parameters. Expression levels from the transcript to the gene level were summarized using tximport R package (Soneson *et al*., 2015) and resulting gene counts were normalized to transcripts per million (TPM). Normalized data was filtered to include only high confidence gene models present in triads, based on the previously published data (Ramírez-González *et al*., 2018). Triads were considered as expressed if they showed expression in at least one homoeolog and an overall triad expression level > 0.5 TPM across the three biological replicates. Any triads with high expression variability between replicates (Euclidean distance > 0.01) were removed from subsequent analysis.

To analyse HEB, a coefficient of variation (CV) was calculated for each triad in every biological replicate as a measure of the relative dispersion of homoeolog TPM around the mean TPM of a triad. Association between HEB and F_5_ line was tested on each triad separately using a linear mixed model approach and analysis of variants (ANOVA) setting genotype as a fixed effect and biological replicate as a random effect as in the following formula:

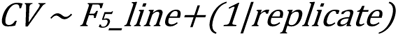

Only triads with highly significant evidence of expression bias variation between F_5_ lines (FDR < 0.0001) were used for further analysis.

To analyse differences in HEB (bias distance) for each triad, the Euclidean distance between the average expression among biological replicates in each F_5_ line and its parental lines was calculated. HEB in F_5_ line was considered significantly divergent from the HEB in parent when the Euclidean distance was 2x the mean bias distance calculated in each F_5_ line population (> 0.2) in > 15 F_5_ lines. These criteria were selected to capture significantly divergent triad expression and account for multiple possible patterns that may be observed for the same triad (i.e. variation between F_5_ individuals). To further discriminate divergent HEB patterns, the following filtering was applied:

- Divergent From One (a) = > 0.2 from Paragon parent & < 0.1 from Charger/Watkins parent in > 15 samples
- Divergent From One (b) = > 0.2 from Charger/Watkins parent & < 0.1 from Paragon parent in > 15 samples
- Divergent From Both = > 0.2 from both parents in > 15 samples
- Conserved = < 0.2 from both parents in > 15 samples

Data manipulation, statistical analysis and image generation were performed using the R language ver. 4.2.1 (R Core Team, 2022) utilizing the lme4 and ggplot2 (Wickham, 2011) packages.

### Gene ontology term enrichment

Gene ontology (GO) was performed as in (Borrill *et al*., 2019). GO terms were transferred from RefSeqv1.0 annotation to the v1.1 annotation for genes which were > 99% identical across > 90% of the sequence. GO enrichment was performed using GOseq R package ver. 1.56.0 (Young *et al*., 2012) separately for each F_5_ line population and bias distance category.

### Variant calling, genotyping and association between genotype and homoeolog expression

Paired RNA-seq reads were quality-trimmed using Trimmomatic ver. 0.39 (Bolger *et al*., 2014) with the following options: LEADING:3 TRAILING:3 SLIDINGWINDOW:4:15 MINLEN:108. They were then mapped to the IWGSC Chinese Spring genome reference v1.0 (IWGSC *et al*., 2018) with HISAT2 ver. 2.1.0 (Kim *et al*., 2019). Using SAMtools ver. 1.12 alignment files for three biological replicates were merged and further processed to exclude unmapped read pairs/mates including reads with mapping quality < 20 (Li *et al*., 2009). Next, PicardTools ver. 2.23.9 (http://broadinstitute.github.io/picard) was used to remove duplicate reads, and finally, variant calling was performed by GATK ver. 4.0.12.0 (McKenna *et al*., 2010) using HaplotypeCaller (Poplin *et al*., 2017) in GVCF mode with subsequent joint genotyping using GenotypeGVCFs. SNP positions were mapped to genes using the RefSeqv1.1 with VariantAnnotation R package (Obenchain *et al*., 2014) and only those that did not overlap with more than one gene were kept. Using in house R script, uniquely mapped SNPs were filtered to keep sites which had < 10% of missing data, read depth ≥ 10 reads in at least five samples, and were homozygous and differed between parental lines. Next, for each F_5_ line, genes which contained more than one SNP and did not display a consistent genotype pattern across the whole sequence (either Paragon, Charger, or Heterozygous) were filtered out to avoid discrepancies in the subsequent analysis. Lastly, F_5_ lines that exhibited unexpectedly high level of heterozygosity (17-36%) were removed from the further analysis.

Using the edgeR package (Robinson *et al*., 2010), raw counts were filtered to include homoeologs with > 0.5 TPM in a triad (see above), normalised by trimmed-mean values and converted to counts per million for further analysis. To test for an association between the homoeolog expression and the inherited genotype, an ANOVA was applied to a linear model based on the following formula:

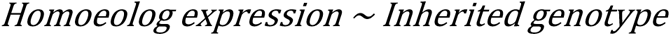

A separate linear model was built for each homoeolog that was expressed and contained genotype calls for > 80% of F_5_ lines. The model was considered significant when the FDR was < 0.05.

Ternary diagrams showing inherited triad expression patterns were plotted using the R package ggtern (Hamilton, 2016).

### SNP genotyping of F_5_ lines for eQTL mapping

eQTL analysis was carried out on 50 F_5_ lines from PxW and 161 F_5_ lines from PxC. The additional 111 PxC F_5_ lines were sequenced in a separate batch as an addition to the 50 used in the first part of our study. A similar strategy as described in the previous section was employed with a few adjustments adequate to the data quality and nature of the analysis. Briefly, only one randomly selected single biological replicate was included in the analysis to ensure a balanced data design in the eQTL analysis. For quality trimming step, the “MINLEN” parameter was reduced from 108 to 60. The alignment files were not merged as only single replicate samples were used for this analysis. All other steps regarding the mapping and variant calling were performed identically to the previous part. A more stringent approach was chosen for filtering of called variants. Using in house R script, uniquely mapped SNPs with a read depth < 3 were labelled as missing data. Sites containing > 2 alleles, > 50% genotype calls missing, displaying > 3% heterozygosity and minor allele frequency < 5% were removed. Lastly, sites that passed this filtering and were homozygous and differed between parental lines were kept for further analysis.

### Quantification of gene expression levels and correction for batch effects

Gene abundance was analysed as previously mentioned using kallisto ver. 0.46.1 (Bray *et al*., 2016) with default parameters and tximpport R package (Soneson *et al*., 2015). A principal component analysis revealed that there was a batch effect separating the data acquired for the previous part of the analysis and the additional data acquired specifically for the eQTL analysis. A correction for the expression levels was applied using combat-seq (Zhang *et al*., 2020) with setting “group” as the population origin (PxC) and “batch” as the batch of samples sequenced together, to remove potential false positive associations detected in the subsequent eQTL discovery analysis.

### Detection of eQTL

The association between SNPs and gene expression was performed by Matrix eQTL R package (Shabalin, 2012) with the setting ‘useModel = modelLINEAR’. The top 20 principal components and phenotyping data for plant height and aerial biomass were used as covariates in the model. All associations with FDR < 0.001 were considered significant. *cis*-eQTLs were defined by location within 1 Mb around the target gene, and *trans*-eQTLs were defined by location on a different chromosome than the target gene or more than 1 Mb from the target gene on the same chromosome. Upset plots displaying the eQTLs were generated with ComplexUpset R package (Krassowski, 2020).

## Results

### HEB is associated with F_5_ line

In order to understand HEB in segregating biparental populations similar to those found within a plant breeding programme, we carried out RNA-seq on two biparental populations at the F_5_ generation. The populations had a common parent Paragon, with the first population Paragon x Charger and the second population Paragon x Watkins landrace WATDE0228. We carried out RNA-seq on 50 F_5_ lines from each population in triplicate, with an average of 43 million reads per sample. We detected expression of 11,215 to 14,331 triads per sample with an average of 13,503 triads in each parental line.

We investigated how these triads were categorised in terms of classic balanced/dominant/suppressed HEB patterns previously described for wheat (Ramírez-González *et al*., 2018). These categories include: balanced with similar expression levels between the three homoeologs, dominant with one homoeolog expressed at a higher level than the other two, and suppressed with a single homoeolog expressed at a lower level than the other two (Figure 1A). After filtering for triad expression, we retained > 12,500 triads (12,717 in PxC; 12,598 in PxW) that display expression level > 0.5 TPM in all 50 F_5_ lines per population. We examined how many triads were uniformly categorised throughout the three biological replicates of every sample. Approximately 25% of triads (2,929 in PxC and 3,179 in PxW) had all three replicates in the same category across all 50 F_5_ lines. Substantial numbers of triads were classified in two categories and triads classified in three categories were occasionally observed (Supplementary Figure 1). The distribution of these discrepancies was highly similar in both populations (Figure 1B). The variation in HEB category between replicates indicates that categorisation may be too stringent and unable to deal with inherent biological variability. Therefore, we instead decided to measure the HEB in F_5_ lines (both PxC and PxW) as the coefficient of variation (CV) between homoeolog expression within each triad (Figure 1A). This quantitative method allowed us to easily identify triads with divergent homoeolog expression, regardless of the precise relative expression category (Figure 1A).

**Figure 1.**
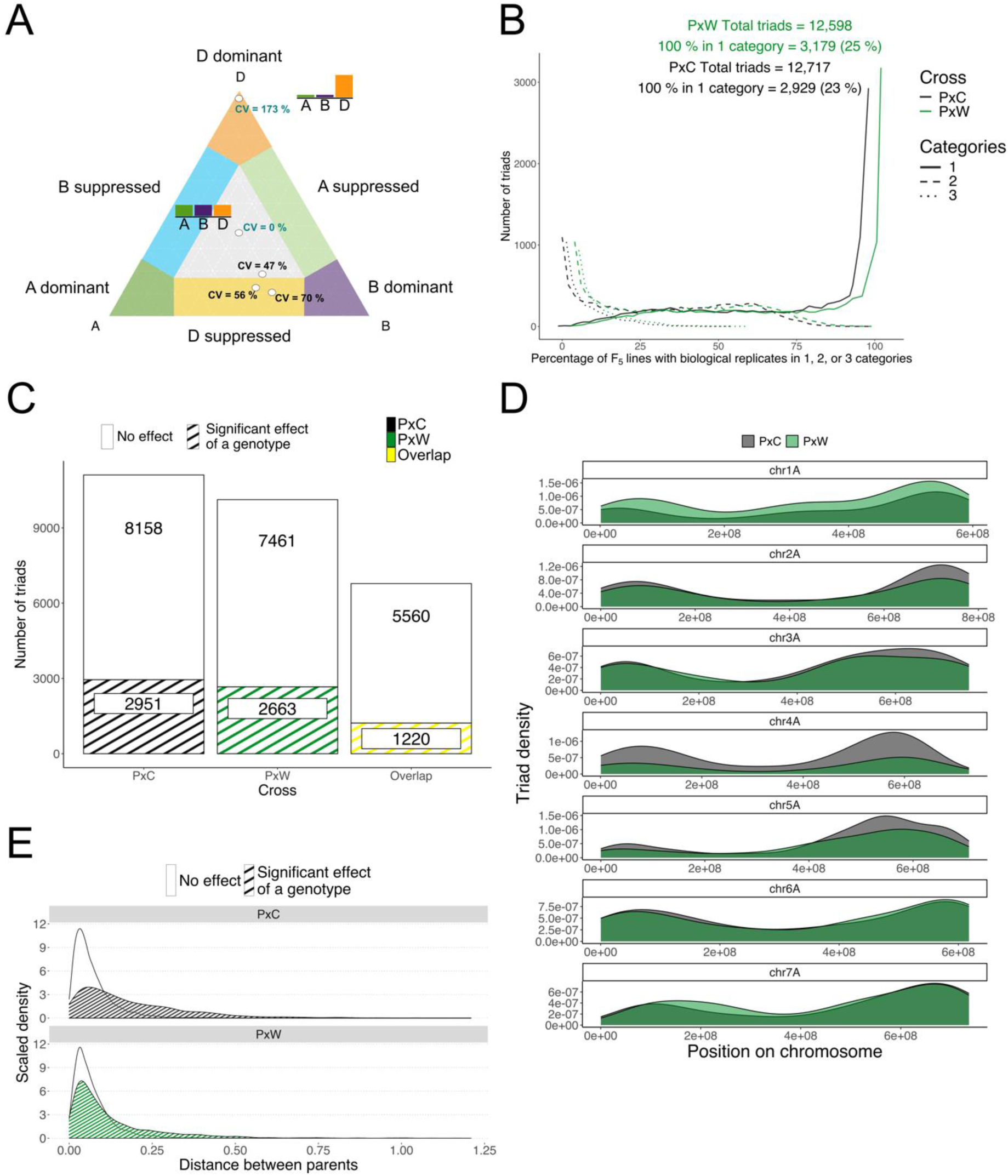
Variation in HEB in F_5_ lines. A) Quantification of HEB in wheat by coefficient of variation (CV) in a triangular plot. Black text represents CV in 3 biological replicates of the same F_5_ line sample. Blue text represents illustrative samples (average of 3 replicates) to show how the CV influences position in the triangular plot with bar plots describing the relative expression of the A, B and D homoeolog in a triad. B) Percentage of F_5_ lines where three biological replicates fall in one (solid line), two (dashed line) or three (dotted line) HEB categories based on their triad expression. Black line represents PxC F_5_ lines, green line represents PxW F_5_ lines. C) Number of triads that showed significant association of HEB (dashed) and those showing no effect (white) in F_5_ lines in the PxC (black) and PxW (green) population, and those which were common to both populations (yellow). D) Triads displaying significant association of HEB to F_5_ line in PxC (black) and PxW (green) population on a subset of chromosomes (1A-7A). E) The effect of HEB divergence between parental lines (Distance between parents) on significant association of HEB in F_5_ line triads in PxC (black) and PxW (green) population (dashed) and those triads showing no effect (white).

Next, we investigated whether the triad expression was linked to the F_5_ line, as that could suggest genetic control occurring at the triad level. After filtering for triad presence in both parental lines for each cross, and using a linear model followed by ANOVA with F_5_ line as a fixed covariate and biological replicate as a random covariate to account for between block microenvironmental differences, we analysed 11,109 and 10,124 triads in PxC and PxW, respectively. We discovered a similar number of triads (2,951 (27%) in PxC and 2,663 (26%) in PxW) displaying association of HEB with F_5_ line (FDR < 0.0001). Moreover, 1,220 of these triads were common for F_5_ lines between both crosses (Figure 1C). We observed differences in the chromosomal position of triads showing association of HEB with F_5_ line between the two populations. A higher number of significant PxC triads were located along chromosome 4 and in distal regions of chromosomes 2, 3 and 5, whereas we detected more significant PxW triads along chromosome 1 and in the proximal region of chromosome 7 (Figure 1D).

We hypothesized that triads which vary significantly in the F_5_ would have a stronger difference in expression bias between parents. Interestingly, the distribution of significant triads was different between populations. In the PxC population, triads with a significant effect of F_5_ line had higher relative expression distances between the parents than non-significant triads suggesting that variable HEB in F_5_ lines were associated with divergent parental HEB (Figure 1E; p < 0.001 Kolmogorov-Smirnov test). In the PxW population the difference between significant and non-significant triads was less pronounced, but still statistically significant (Figure 1E; p < 0.001 Kolmogorov-Smirnov test). However, in both populations, many of the significant triads (68-74%) had a low bias distance between the parents (< 0.1) indicating that HEB in F_5_ lines could occur *de novo*, despite similar HEB in the parents.

### HEB is associated with inherited genotype

To understand heritable effects on HEB, we focused our subsequent analysis only on triads for which F_5_ line had a significant effect (2,951 triads in PxC and 2,663 triads in PxW) and calculated the expression bias distance between each F_5_ line and their parents. The majority of triads in each population displayed bias distance < 0.2, indicating that in the F_5_ lines the triads were similar to one or both parents in their HEB (Figure 2A, B). We discovered that triads can display distinctive patterns of biased expression in the F_5_ lines (Figure 2C), which we categorised according to the bias distance to each parent (coloured sectors on Figure 2A and B). We considered triads which had a bias distance < 0.2 from both parents as conserved i.e. HEB does not differ strongly between parents and F_5_ lines. Triads displaying > 0.2 bias distance from one parent and < 0.1 from the other parent were considered to be divergent from one parent (DFO), whether this was divergent from Paragon parent (DFO_a) or divergent from Charger/Watkins parent (DFO_b). Triads with a bias distance > 0.2 from both parents were considered divergent from both parents (DFB). We investigated how these patterns varied inside each population by calculating the percentage of F_5_ lines showing a specific pattern. Whereas triads showing the conserved pattern seemed to be highly uniform across F_5_ lines in both populations, other triads varied in their pattern between F_5_ lines (Figure 2D and E). This was highly noticeable in DFO patterns where a portion of triads exhibited divergence from one parent and another portion exhibited divergence from the other parent (Figure 2D, E). These DFO triads with opposite divergences in individual F_5_ lines were more frequent in the PxC population than in PxW (Figure 2D), whilst triads which diverged from both parents were more common in the PxW population (Figure 2E). In order to quantify these triads based on their patterns more precisely, we implemented a condition of at least 15 F_5_ lines out of 50 (30%) showing a specific pattern. This allowed us to classify 2,814 triads in PxC and 2,573 triads in PxW (out of the total 2,951 triads in PC and 2,663 triads in PW). Setting our thresholds this way left a few triads unclassified, however this was < 5% of the total number of triads.

**Figure 2.**
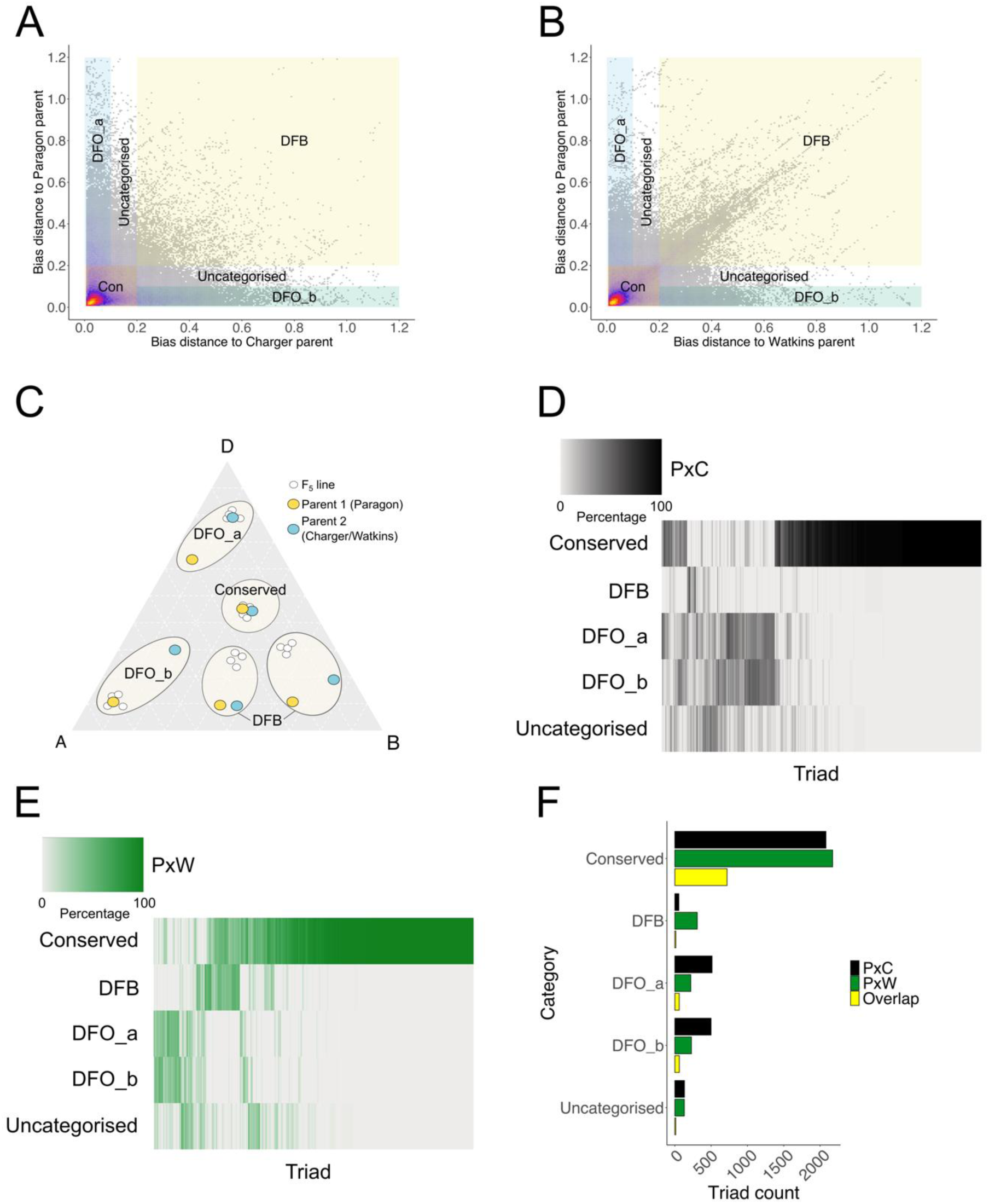
HEB profiles in parental lines influence the HEB in F_5._ A) and B) Differences between HEB in F_5_ lines and HEB in Paragon and Charger parental lines (A), or Paragon and Watkins parental lines (B). Areas are classified based on the expression bias distance calculated between F_5_ lines and parental lines: DFO_a – divergent from the Paragon parent, DFO_b – divergent from the Watkins or Charger parent, DFB – divergent from both parents, Con – conserved, and Uncategorised. C) Examples of distinctive patterns of biased expression between F_5_ lines (white points) and their parents (Orange – Paragon parent, Blue – Charger/Watkins parent). D) and E) Variability in triad expression patterns in PxC (D) and PxW (E) F_5_ lines. Scale represents the percentage of F_5_ lines showing the same pattern from 0% (grey) to 100% (black – PxC; green - PxW). F) Number of triads exhibiting a specific expression pattern as compared to parental lines based on bias distance calculations for PxC (black) and PxW (green) F_5_ lines and an overlap between those two populations (yellow).

The majority of triads showed a conserved pattern (2,081 triads in PxC and 2,173 in PxW), and 33% of these triads were shared between both populations (719 triads). There was no significant difference in the number of triads displaying divergence from one parent or the other (DFO_a and DFO_b) in each F_5_ population (513 and 499 in PxC; 220 and 229 in PxW) indicating no parental bias (Figure 2F). Despite both populations containing a similar number of triads which were affected by genotype (2,951 in PxC vs 2,663 in PxW), we detected more than twice the number of triads showing divergence from one parent (but similar to the other parent) in PxC compared to PxW F_5_ population (Figure 2F). This suggests that when there is a greater bias distance separating the parents (Figure 1E), the offspring tend to resemble the value of one of the parents rather than an average. By further examination we discovered that there was a high overlap in triads showing DFO_a or DFO_b pattern (330 in PxC Figure 2D, F; 155 in PxW Figure 2E, F), meaning that these particular triads highly follow the expression of either one, or the other parent in F_5_ lines. Moreover, this happens in a highly balanced way with an average of around 20 out of 50 F_5_ lines following the expression of one or the other parent (Supplementary Figure 2A). This nearly 1:1 segregation may reflect a direct association between the HEB and an inherited parental genotype.

Interestingly, there were significantly more triads displaying DFB pattern in PxW (309) than PxC (56) (Figure 2F; also visible as a diagonal cluster of dots in Figure 2B). In PxC F_5_ lines, the majority of DFB triads (88%) displayed a high divergence in HEB between the parents (bias distance between parental lines > 0.2). However, in PxW F_5_ lines, it was the opposite where the majority of DFB triads (84%) displayed low initial divergence in HEB (bias distance between parental lines < 0.2) (Supplementary Figure 2B). This similarity in parental expression for DFB triads in PxW F_5_ lines could either point to non-genetic factors influencing the HEB or a specific combination of inherited parental homoeologs that ultimately lead to a novel HEB in F_5_ lines. Few triads were identified as divergent from one or both parents in both populations suggesting that divergent HEB in the F_5_ were largely driven by population specific factors (94 triads; Figure 2F).

We investigated whether these patterns of biased expression in the F_5_ lines correlated with the function of the triads. Triads in all bias categories and both populations were enriched for gene ontology terms associated with four major core plant cell function areas: photosynthesis, glucose metabolism, transcription and translation (Supplementary Table 1). There were small population specific effects: for example, GO terms related to photosynthesis were more strongly related to triads which followed the expression of one of the two parents (DFO_a or DFO_b) in PxC, whereas in PxW associations with photosynthesis were stronger for triads which significantly differed in their expression compared to both parents (DFB) (Supplementary Table 1). This hints that core cell processes may be affected differently in various populations.

### Within homoeolog SNPs explain variation in HEB

To test the direct association between HEB and inherited genotype, we performed variant calling resulting in 58,845 and 29,262 SNPs between PxC and PxW parental lines, respectively. We filtered homoeolog genotype information in F_5_ lines based on several criteria including removing homoeologs which contained > 1 SNP in their sequence that did not provide a consistent genotype. From the ∼2,500 triads which had significant HEB associated with genotype in each population, we were able to identify reliable SNPs in 1,591 homoeologs in PxC and 703 homoeologs in PxW, corresponding to 1,306 triads in PxC and 658 triads in PxW (Supplementary Tables 2 and 3). Using a linear model followed by ANOVA we found that the homoeolog expression was associated with inherited genotype in 69% of these homoeologs in each population, corresponding to 980 triads in PxC and 464 triads in PxW (adj. p < 0.05).

We investigated whether heritable changes to HEB were associated with one or more homoeologs. In PxC, out of 308 triads displaying bias distance > 0.2 between the Paragon and Charger parent, 89.3% had a significant difference in HEB in the F_5_ lines which was associated with a change in the expression of one homoeolog (Table 1). In PxW, we observed the same trend with 93.3% of 119 divergent triads which were associated with change in gene expression and genotype in one homoeolog (Table 1). In both populations a small number of triads had two homoeologs associated with a change in expression (10.4% of divergent triads in PxC; 6.7% of divergent triads in PxW).

**Table 1.** Association of homoeolog expression with inherited genotype in F_5_ populations in triads where bias distance is > 0.2 between parents.

| **Homoeologs(s) with genotype associated with gene expression** | **PxC (# of triads)** | **PxW (# of triads)** |
| --- | --- | --- |
| A | 115 | 57 |
| B | 116 | 36 |
| D | 44 | 18 |
| **Total with one homoeolog associated** | **275 (89.3%)** | **111 (93.3%)** |
| A and B | 17 | 4 |
| A and D | 9 | 2 |
| B and D | 6 | 2 |
| **Total with two homoeologs associated** | **32 (10.4%)** | **8 (6.7%)** |
| A and B and D | 1 (0.32%) | 0 |
| **Total with one, two or three homoeologs associated** | **308** | **119** |

In the majority of triads (60% in PxC and 62% in PxW), the homoeolog(s) that were found to have an association between the inherited genotype and expression had higher variation in normalised expression values compared to the other homoeologs in the triad (Figure 3A). Therefore, a change in the expression of one homoeolog correlates with the bias distance calculations and can discriminate F_5_ lines where the HEB of a respective triad is divergent from one or the other parent (Figure 3B). These subpopulations of F_5_ lines can be discriminated based on their inherited genotype by the relative homoeolog expression levels on a triangular plot, with the variably expressed homoeolog driving the separation between subpopulations (Figure 3C). After further inspection the triads that did not follow this pattern, we discovered 14% and 12% triads in PxC and PxW, respectively, where the homoeolog with the highest variation in expression did not show association between inherited genotype and expression, which was frequently due to incomplete genotype information for this highly variable homoeolog. Lastly, 26% of triads in both populations showed the highest variation in the expression of a homoeolog without any genotype information based on our RNA-seq data.

**Figure 3.**
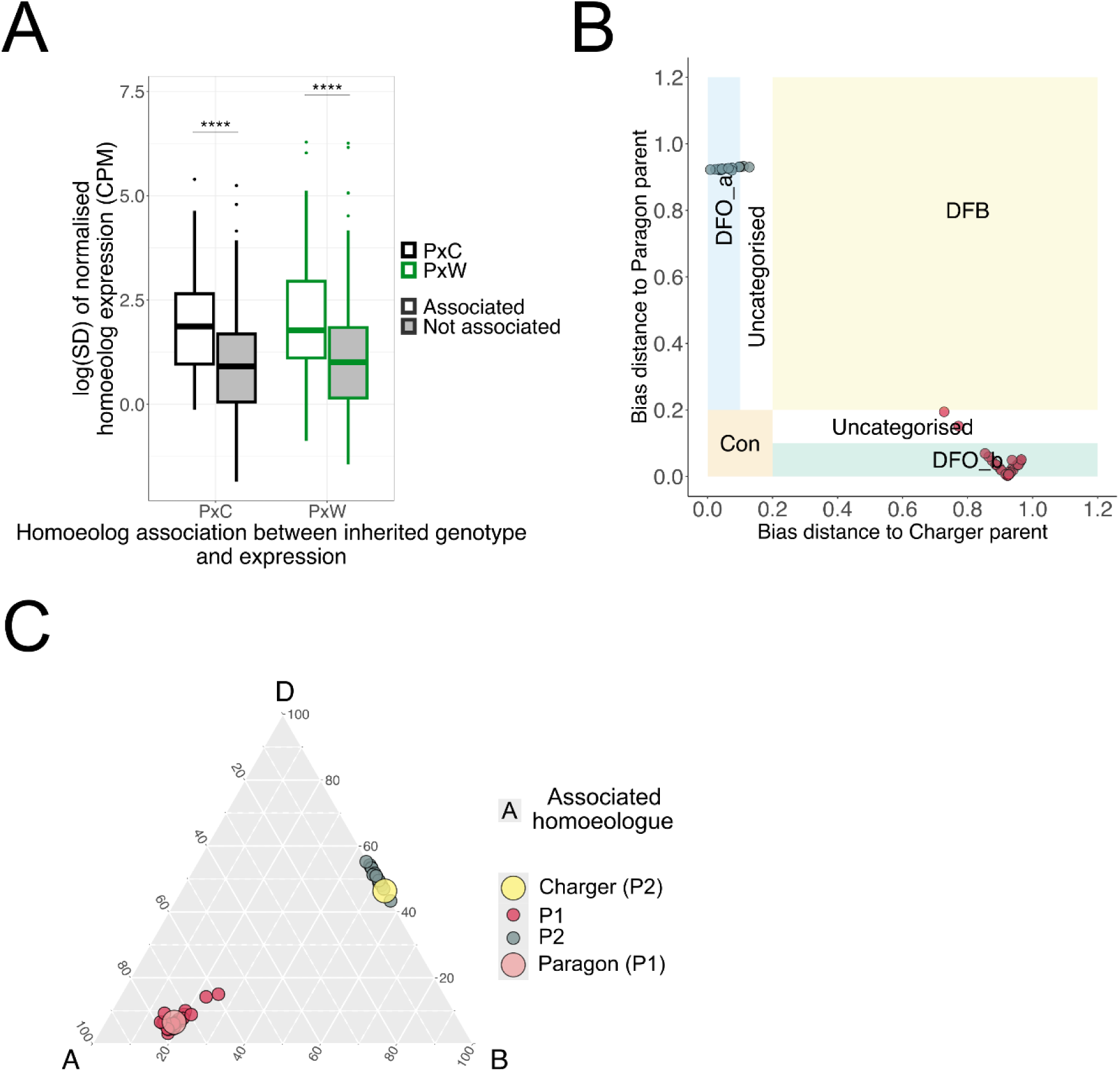
HEB is associated with inherited genotype. A) Variation in the expression of homoeologs that were found to have an association between expression and inherited genotype (Associated - white), and in the expression of other two homoeologs in the same triad (Not associated - grey) in two F_5_ populations (PxC – black, PxW – green). Variation is represented as logarithm of standard deviation (SD). Asterisks represent the significant difference in variation between associated and not associated homoeologs (Wilcoxon rank sum test; **** p < 0.0001). B) The distribution of PxC F_5_ lines in an example triad (*TraesCS1A02G004200*, *TraesCS1B02G005600*, *TraesCS1D02G003300*) displaying association between the triad expression and inherited genotype. The F_5_ lines are either similar to the Paragon or Charger parent based on the bias distance calculated in each sample towards the parental lines. Colours of dots represent the genotype inherited (Paragon inherited genotype = red; Charger inherited genotype = grey). C) HEB differences between F_5_ lines for the same example triad (*TraesCS1A02G004200*, *TraesCS1B02G005600*, *TraesCS1D02G003300)* show that the association between the triad expression and inherited genotype is driven by A homoeolog. The position of every F_5_ line and the parental samples in the triangle is calculated from the TPM expression values. The size of dots differentiates parental samples from F_5_ samples. Colour of dots represents the genotype of each sample (Paragon parent = pink; Charger parent = yellow; Paragon inherited genotype = red; Charger inherited genotype = grey).

### Changes to expression level are associated with *cis*-variants

To further confirm whether there is a direct connection between inherited genotype and expression, and discover loci that might contribute to this inheritance, we conducted an eQTL analysis. To carry out the analysis with balanced sampling, we randomly selected 1 replicate from the 3 biological replicates previously sequenced. To increase the statistical power, for the PxC F_5_ line population, we isolated and sequenced RNA from additional F_5_ lines which were harvested at the same time as the material in the previous part. In total, we used 161 PxC and 50 PxW F_5_ lines for eQTL analysis. We applied multiple stringent filters to remove low-quality regions (see Methods) and in total, we retrieved 7,957 and 5,538 biallelic SNPs from 3,080 and 2,059 genes for PxC and PxW F_5_ lines, respectively (Figure 4A).

**Figure 4.**
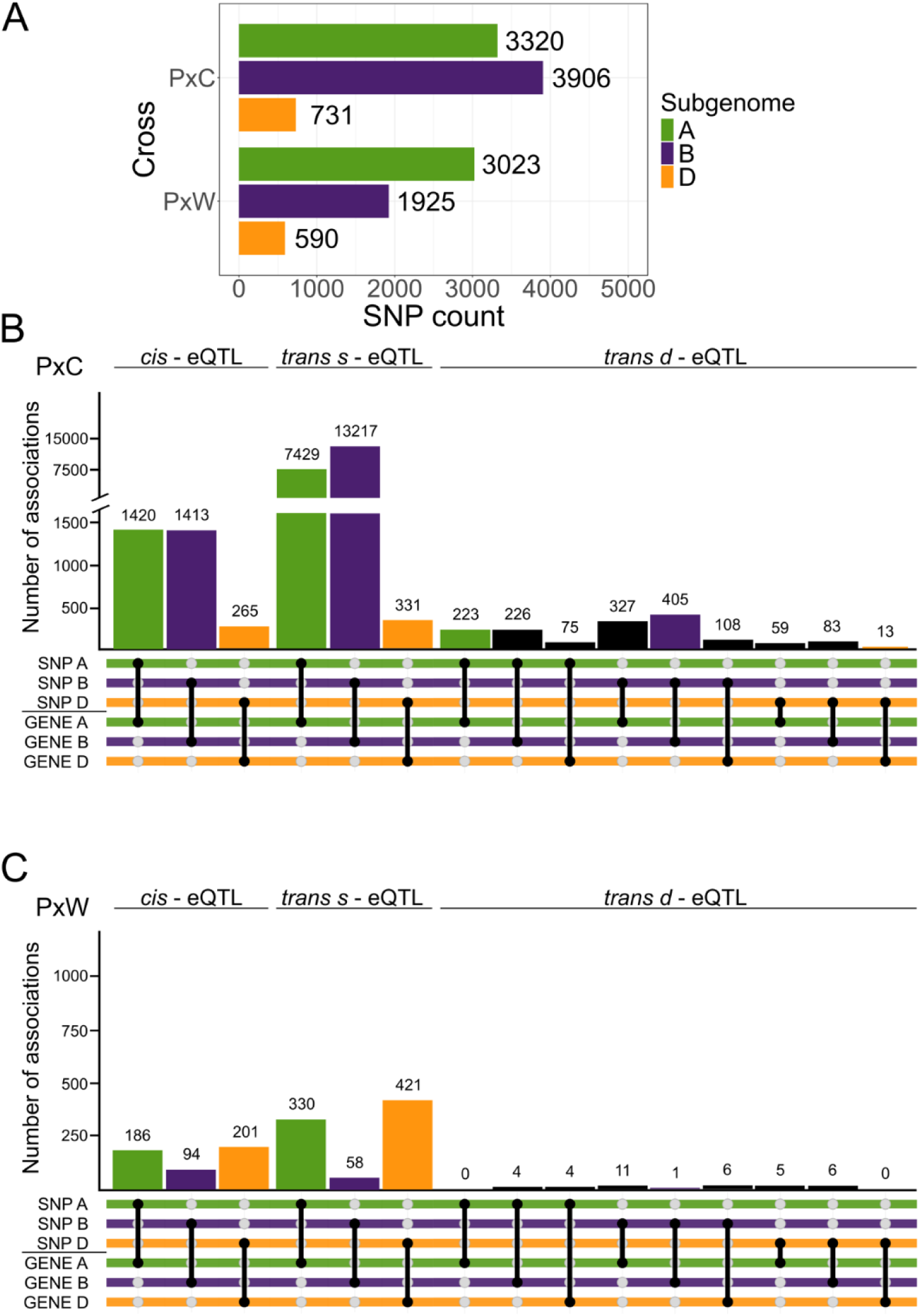
eQTL analysis confirms association between homoeolog expression and inherited genotype. A) Number of SNPs identified between parental lines in each population. Colours correspond with the specific subgenome (green = A; purple = B; orange = D). B) and C) Number of associations between SNPs and homoeolog expression identified using eQTL analysis in B) PxC and C) PxW population. eQTL types are defined as *cis-* (same chromosome < 1Mb); *trans-s-* (same chromosome > 1Mb) and *trans-d-* (different chromosome) eQTLs. Coloured bars represent direct association between a SNP and gene expression in the same subgenome. Black bars represent direct association between a SNP and gene expression in different subgenomes.

We identified a significantly higher total number of eQTLs (FDR < 0.001) in PxC (55,484) than PxW (3,064). Although the number of high-quality SNPs was similar between both F_5_ line populations, the low number of input samples in PxW F_5_ lines might have led to reduction of statistical power needed for the identification of eQTLs. In both populations, the majority of eQTLs identified were *trans-*same-eQTLs (*trans-*s-eQTLs) where the SNP and associated expressed gene were on the same chromosome but > 1Mb apart, representing within chromosome long-range interactions and/or associations (Figure 4B and C). *Cis*-eQTLs, where the distance between the SNP and the associated expressed gene was < 1Mb on the same chromosome, formed the second most abundant group. There was a higher number of *cis*- and *trans*-s-eQTLs identified in A and B subgenomes than the D subgenome in the PxC population. In the PxW population, a higher number of *cis*- and *trans*-s-eQTLs were identified in A and D subgenome than the B subgenome. These results show a high number of SNPs are associated with expression of different gene/s or clusters along the same chromosome, consistent with our findings above that HEB is associated with the inherited genotype at that locus (or linked loci).

We also discovered a higher number of *trans-*different-eQTLs (*trans*-d-eQTLs), where the SNP and associated expressed gene were on different chromosomes regardless of sub-genome, in the PxC population compared to the PxW population (Figure 4B and C). The PxC population A and B subgenomes had more associations between SNPs and gene expression on their own subgenome, and between each other, as compared to associations with the D subgenome. Moreover, the D subgenome also displayed this pattern with the D SNPs having more associations with genes expressed from A and B subgenome, rather than with genes expressed on D subgenome, in both populations. The latter could be attributed to the fact, that we have identified about five times fewer SNPs on the D subgenome in general.

Lastly, we examined whether the eQTLs we discovered might be associated with previously identified agriculturally relevant traits. Based on data from (Cheng *et al*., 2024), we were able to identify a number of loci in PxC which correspond to the physical marker position for several traits, such as grain width, heading time and plant height (Supplementary Table 4). This suggests that differences in HEB between cultivars is associated with variation at agriculturally-relevant loci.

## Discussion

In this work, we examined what happens to HEB over short time periods (five generations) which could be representative of wheat breeding programmes. We found that in two biparental mapping populations, homoeolog expression bias is inherited for 26-27% of triads in the F_5_ generation. Most of these triads (∼ 70%) have a conserved expression similar to the parents, however novel patterns of HEB were also generated, even if parents had highly similar HEB. This occurred more commonly in the elite x landrace population, than in the population with two elite parents. We discovered that inherited HEB patterns were largely driven by changes in the expression of one homoeolog (∼ 60%), allowing HEB in subsequent generations to match parental expression HEB. We also found that HEB is altered by both *cis-* and *trans-*variants altering expression within the subgenome and some of the triads affected may be related to agriculturally important traits.

### HEB is inherited for over a quarter of triads and influenced mainly by a single homoeolog

Within species variation in HEB, and how this persists between generations after crossing is not well understood, yet could have significant implications for allopolyploid crop breeding. Here, we found that HEB is inherited for 26-27% of triads in biparental hexaploid wheat populations. The triads which do not show heritable HEB are frequently invariant between parents. It is likely that different triads would demonstrate heritable changes in HEB in other mapping populations since wheat cultivars demonstrate variable HEB (He *et al*., 2022; Wang *et al*., 2024).

We found that the majority of triads have a conserved expression similar to the parents, however novel patterns of HEB were also generated, even if parents had highly similar HEB, indicating transgressive patterns of HEB within the progeny. We found that a substantial proportion of triads segregate for HEB in the F_5_ generation, with individual F_5_ progeny showing a similar HEB to one parent or the other (Figure 5A and B). This binary divergence between HEB is consistent with the strong influence of *cis*-regulation which we detected within these populations and was identified as regulating HEB previously in wheat (He *et al*., 2022). It is likely that the divergence between HEB patterns after crossing is rapidly established within each progeny because even in a more extreme scenario of resynthesizing allopolyploid *Brassica napus*, HEB is established within two generations and persisted across all eight generations tested (Bird *et al*., 2021).

**Figure 5.**
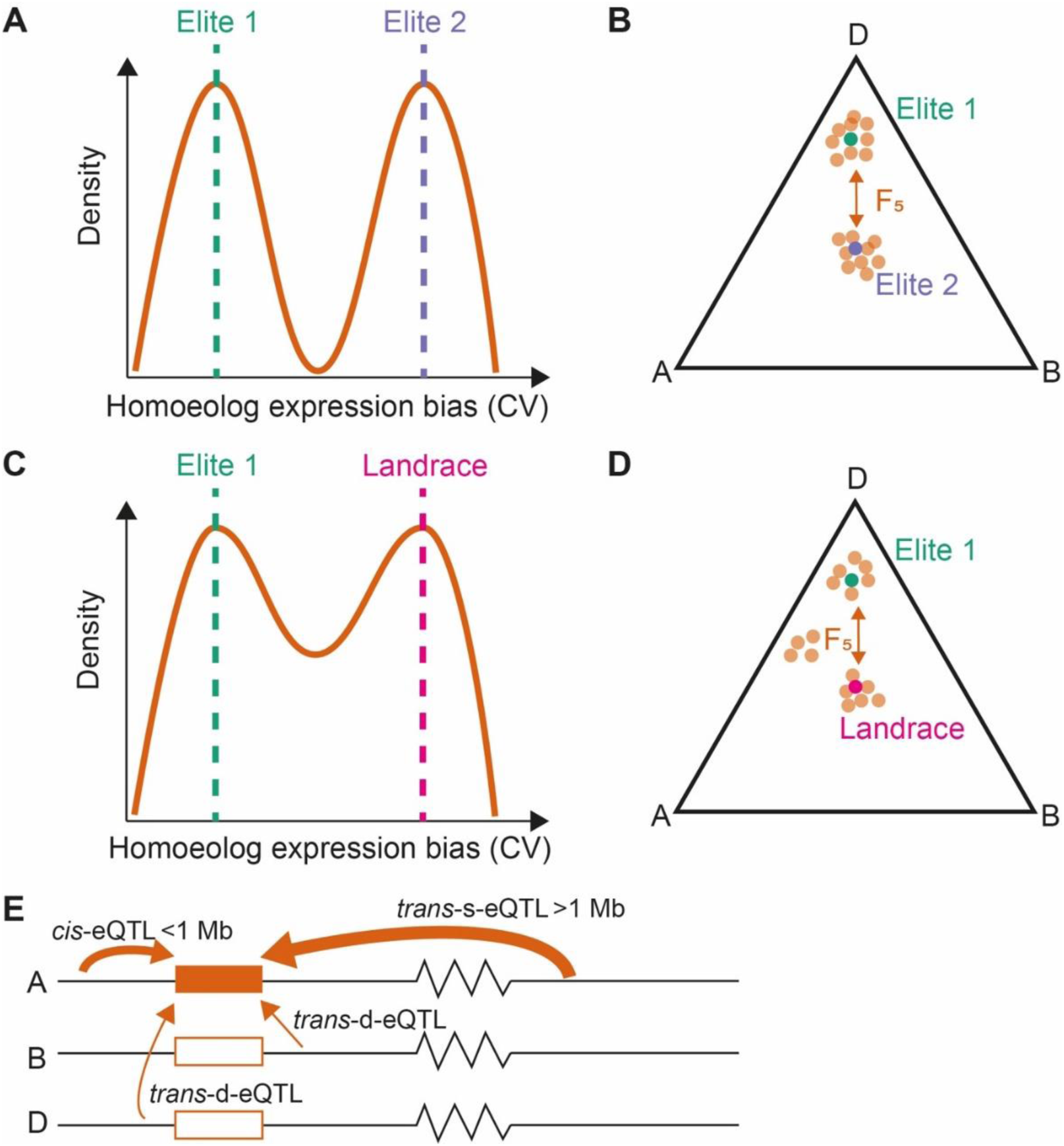
Summary of homoeolog expression bias (HEB) in the F_5_ generation of biparental crosses. In an elite x elite cross (A and B), the coefficient of variation (CV) between homoeologs in F_5_ individuals tends to match either one parent or the other, represented as a density line graph (A) and within a triangle plot (B). In an elite x landrace cross (C and D) the CV between homoeologs in F_5_ individuals matched either one parent or the other, or displayed a pattern different from both parents, represented as density line graph (C) and within a triangle plot (D). E) HEB is mainly influenced by within chromosome eQTL which are mostly > 1Mb away from the gene (*trans*-s-eQTL), *cis*-eQTL also have a substantial role, while interactions between chromosomes through *trans*-d-eQTL are rarer. The width of the arrows represents the number of eQTL.

Individual triads behave differently in the two different populations, although > 40% of triads which are heritable in HEB were common in both populations which may be caused by the shared Paragon parent, or suggest specific properties of these triads. We found that these common triads were evenly distributed along all chromosomes, whereas a number of triads unique to each population showed asymmetrical distribution on several chromosomes. Minor differences in HEB patterns in progeny of crosses with one common parent were previously observed in two synthesized *B. napus* F_1_ hybrids. Whereas A_n_C_o_ hybrid displayed higher number of C-biased homoeologs, A_r_C_o_ hybrid displayed no significant difference between the number of A- and C-biased homoeologs (Zhang *et al*., 2018). In our study, a substantial proportion of triads in the PxC population switched between parental HEB in F_5_ lines, which was largely driven by changes in the expression of one homeolog. However, a larger number of triads in the PxW population were divergent from both parents (DFB) and these novel HEB patterns could not be explained by genotype information retrieved from coding sequences, suggesting that the causative factors of this behaviour might be differences in regulatory sequences or/and epigenetic modifications which have been previously linked to HEB (Ramírez-González *et al*., 2018; Li *et al*., 2021), or missing genotype information from some samples due to low sequence depth. This establishment of novel HEB patterns (DFB) occurred more commonly in the PxW population which had a landrace parent, than in the population between elite cultivars (PxC) (Figure 5C and D).

### *Cis*-regulation and same chromosome *trans* interactions are a major driver of HEB

HEB in wheat is explained by a combination of both *cis-* and *trans-*acting variants, with the majority of these coming from the A and B subgenomes which are more genetically diverse than the D subgenome. In the context of genome organisation, it was previously demonstrated that wheat chromatin architecture is not random and the A and B subgenomes interact most frequently with themselves and more frequently with each other than with the D subgenome, although the D subgenome displays higher chromatin accessibility than the A and B subgenomes due to the lower transposable element content (Concia *et al*., 2020; Jordan *et al*., 2020). This agrees with our results in the PxC population where *trans-*s- and *cis-*eQTLs (i.e. within chromosome associations) identified in the A and B subgenomes were the most abundant (Figure 5E), and *trans-*d-eQTL numbers were more comparable between A and B subgenomes. Due to our experimental design using biparental populations, the number of *trans-*s-variants discovered is mainly correlative and describes stable patterns of inherited HEB rather than regulatory relationships. However, the importance of within-chromosome regulation of HEB is confirmed by recent work comparing accessions, which reduces this limitation (Wang *et al*., 2024).

A limitation of our current work and studies of HEB between wheat accessions (He *et al*., 2022; Wang *et al*., 2024) is the use of a single reference genome. The introduction of a pantranscriptome approach (Coombes *et al*., 2024) has been shown to improve the accuracy of quantification for 4-9% of genes which are underestimated using a single reference. Therefore, in future studies, the use of a pantranscriptome approach, or even tailored reference sequences will be valuable for correct identification and description of HEB dynamics in different crop varieties. Nevertheless, our study which uses biparental populations, suffers less from this source of bias, than studies which use hundreds of accessions which may be more highly impacted by genetic diversity.

### Implications for breeding allopolyploids

Much current wheat research focuses on improving elite cultivars by introducing novel alleles from wild relatives of wheat and unexplored landrace cultivars to combat challenges such as low bioavailability of minerals, new strains of pathogens and other challenges emerging due to climate change (Wingen *et al*., 2014; Cruppe *et al*., 2020; Fatima *et al*., 2020; Pour-Aboughadareh *et al*., 2021). We found that novel HEB patterns were generated more frequently in the population containing a landrace. This raises a new challenge for wheat researchers as we increasingly introduce additional genetic diversity from outside the conventional elite germplasm - we may be generating unpredictable HEB in triads which may affect the resulting phenotype for a desired trait. This result was obtained by following a traditional crossing scheme and therefore, in the future, it might be preferable to focus on creating novel allelic variants or mimicking natural variants using precision breeding techniques such as CRISPR to avoid unintended consequences (Gao *et al*., 2024).

### Updated approach to analyse HEB in high level polyploids

Here we developed a new framework beyond HEB categories to analyse HEB in hexaploid wheat (Ramírez-González *et al*., 2018). We found that the variation in HEB between biological replicates causes problems in accurate categorisation. Therefore, we developed an alternative approach based on calculating the coefficient of variation in homoeolog expression which allowed us to properly test the effect of genotype on inherited triad expression in a great number of triads. We suggest that this approach may be more appropriate to study HEB in higher level allopolyploids, especially in those containing more than 2 subgenomes, such as wheat (3 subgenomes), oat (3 subgenomes) or strawberry where the HEB has to be calculated in a three-dimensional tetrahedron space due to the 4 subgenomes present (Jin *et al*., 2023).

In conclusion, our results indicate that there is significant reprogramming and stabilisation of HEB within five generations that differs significantly based on the parental lines used in the crossing. The influence of these changes on phenotype and impact on allopolyploid crop breeding requires further investigation.

## Acknowledgements

We thank Richard Morris for insightful comments and feedback on the manuscript, Luzie Wingen for providing access to phenotypic data and Catherine Evans and Muhammad Waqas Ali for help with harvesting the plant material. The research was funded by the UK Biotechnology and Biological Sciences Research Council (BBSRC) through grant BB/T013524/2 and the Institute Strategic Programmes Delivering Sustainable Wheat (DSW) (BB/X011003/1) and Building Robustness in Crops (BRiC) (BB/X01102X/1). RA was funded by a fellowship from the Peter and Traudl Engelhorn Foundation. This study was supported by the NBI Research Computing group through HPC resources.

## Author contributions

PB designed the study. PB, RA, SB and SLM collected the data. MG and RA analysed the data with assistance from SB. XW provided intellectual input. MG and PB wrote the initial draft of the manuscript. All authors reviewed, edited the manuscript and gave approval for submission of the final version. MG and RA contributed equally to this work.

## Data availability

The data that supports the findings of this study are available in public repositories. Raw RNA-seq data can be obtained through BioProject ID PRJNA1128551 on the NCBI Sequence Read Archive. Scripts and intermediate data tables necessary to run them and create plots for figures are available on GitHub (https://github.com/Borrill-Lab/Inheritance-of-HEB-in-wheat) or figshare (https://figshare.com/projects/Reprogramming_and_stabilisation_of_homoeolog_expression_bias_in_hexaploid_wheat_biparental_populations/214495) depending on their size.

## Supporting Information

**Supplementary Figure 1.** Triad expression patterns in two representative samples according to their classification into multiple expression categories.

**Supplementary Figure 2.** Bias distance between HEB in PxC and PxW F_6_ lines and their parents.

**Supplementary Table 1.** Gene Ontology enrichment of triads in F_5_ lines in groups classified by calculating bias distance between F_5_ lines and parental lines.

**Supplementary Table 2.** Summary of triad expression data in PxC F_5_ lines, homoeologs associated with change in triad expression and bias distance calculated between F_5_ lines and parental lines.

**Supplementary Table 3.** Summary of triad expression data in PxW F_5_ lines, homoeologs associated with change in triad expression and bias distance calculated between F_5_ lines and parental lines.

**Supplementary Table 4.** Selected eQTLs identified in the PxC population, their association to specific traits and identified rice orthologs.

