## Supplementary Figures and Supplementary Table 4 for "Rapid reprogramming and stabilisation of homoeolog expression bias in hexaploid wheat biparental populations"

**Supplementary Figures and Tables for "Rapid reprogramming and stabilisation of homoeolog expression bias in hexaploid wheat biparental populations".**

Marek Glombik[1]*, Ramesh Arunkumar[1]*[#], Samuel Burrows[1], Sophie Louise Mogg[2§], Xiaoming Wang[3], Philippa Borrill[1†]

[1] Department of Crop Genetics, John Innes Centre, Norwich Research Park, Norwich, NR4 7UH, UK.

[2] School of Biosciences, University of Birmingham, Birmingham, B15 2TT, UK.

[3] State Key Laboratory for Crop Stress Resistance and High-Efficiency Production, College of Agronomy, Northwest A&F University, Yangling, Shaanxi, China.

* These authors contributed equally.

[#] Present address: School of Life Sciences, Technical University of Munich, Alte Akademie 8, 85354 Freising, Germany.

§ Present address: School of Biological Sciences, University of Manchester, Manchester, M13 9PL, UK.

### Supplementary Figures

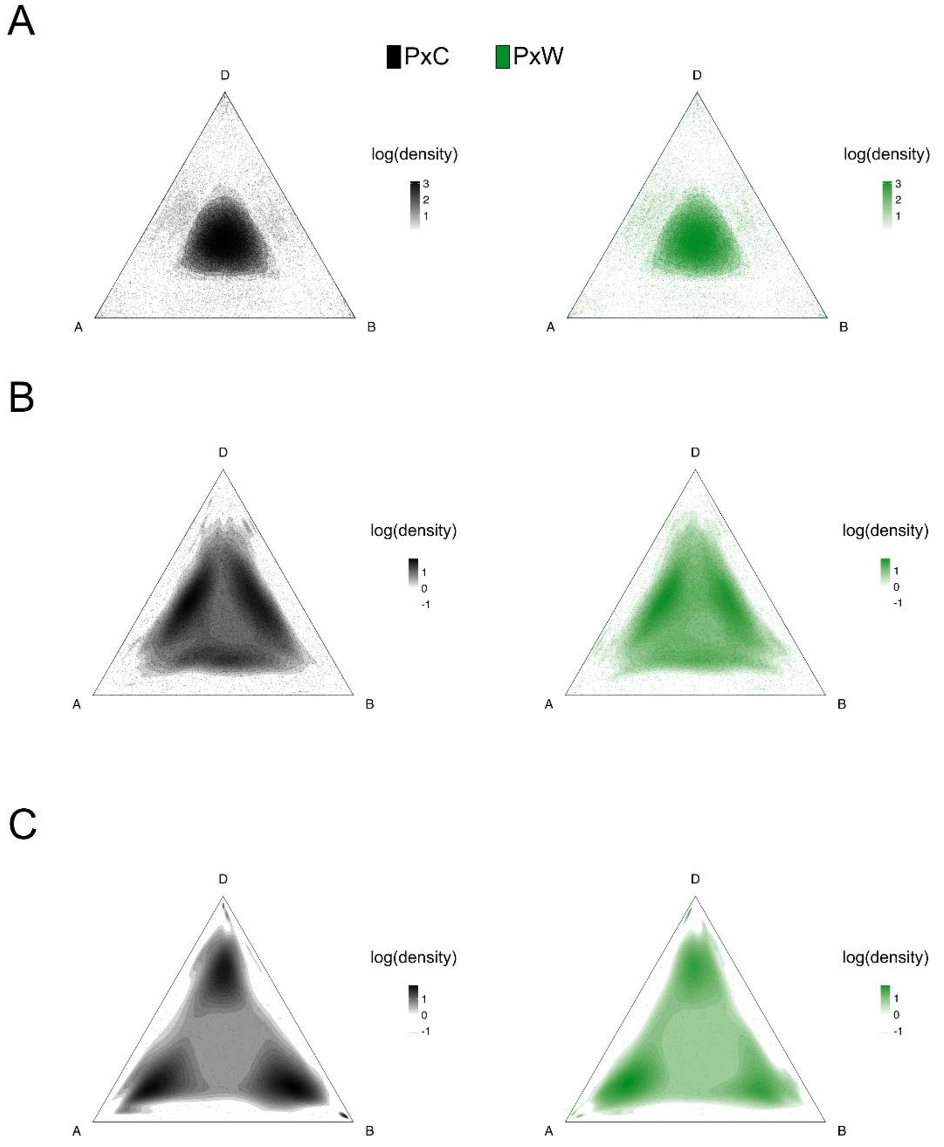


**Supplementary Figure 1.** Triad expression patterns in two representative samples according to their classification into multiple expression categories. A-C) Expression pattern of triads having all three biological replicates in one, two, or three categories, respectively. PxC $F_5$ representative sample (black); PxW $F_5$ representative sample (green).

A

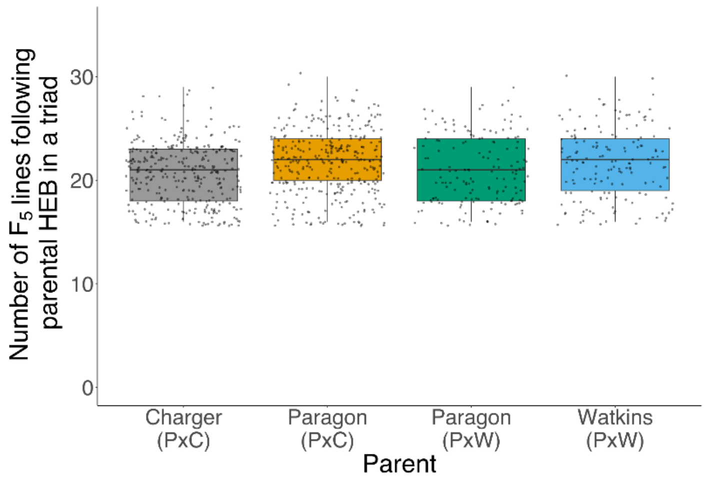


B

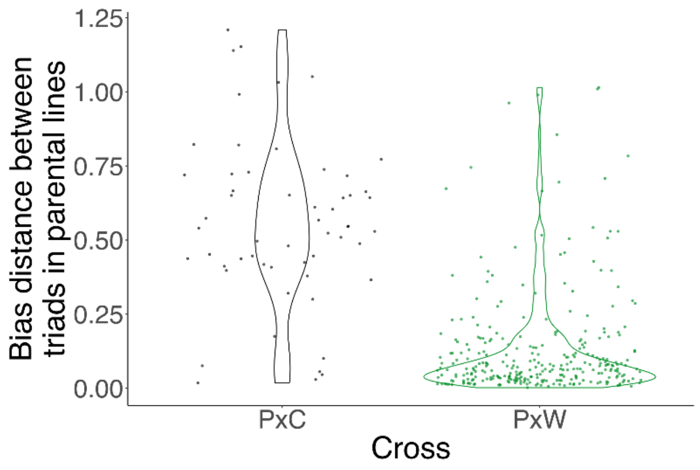


**Supplementary Figure 2.** Bias distance between HEB in PxC and PxW $F_6$ lines and their parents. A) The number of $F_6$ lines which followed the expression pattern of each parent for triads which overlapped between DFO_a and DFO_b (330 in PxC; 115 in PxW). B) The bias distance between parents for triads which were different from both parents (DFB) in the $F_6$ generation.

### Supplementary Tables

**Supplementary Table 1.** Gene Ontology enrichment of triads in $F_5$ lines in groups classified by calculating bias distance between $F_5$ lines and parental lines.

See separate excel file.

**Supplementary Table 2.** Summary of triad expression data in PxC $F_5$ lines, homoeologs associated with change in triad expression and bias distance calculated between $F_5$ lines and parental lines.

See separate excel file.

**Supplementary Table 3.** Summary of triad expression data in PxW $F_5$ lines, homoeologs associated with change in triad expression and bias distance calculated between $F_5$ lines and parental lines.

See separate excel file.

**Supplementary Table 4.** Selected eQTLs identified in the PxC population, their association to specific traits and identified rice orthologs.

| SNP from | Associated gene expression | Function | eQTL type | Associated trait in PxC | Associated trait in other studies | Source |
|---|---|---|---|---|---|---|
| *TraesCS4A02G382400* | | TaHDZ22-4A; Homeobox domain-containing protein | *cis-* | *Heading* | response to stress | https://doi.org/10.1186/s12870-020-2252-6 |
| | *TraesCS4A02G382900* | RING-type E3 ubiquitin transferase | | | stripe rust resistance | https://doi.org/10.1186/s12864-023-09336-y |
| *TraesCS4A02G298600* | | HSP20-like chaperone | *cis-* | *Grain width* | heat stress | https://doi.org/10.3390/plants11212987 |
| | *TraesCS4A02G299200* | Co-chaperone protein p23 | | | NA | NA |
| *TraesCS4A02G461700* | | NB-ARC domain-containing protein | *trans same -* | *Plant height* | NA | NA |
| | *TraesCS4A02G499900* | NAC domain-containing protein | | | heat stress tolerance | https://doi.org/10.1111/pbi.13529 |
